# Superposition of target structures enables design of bi-stable RNA molecules with deep learning

**DOI:** 10.1101/2023.03.11.532170

**Authors:** Christopher F. Blum, Petra Kolkhof, Markus Kollmann

**Affiliations:** Institute for Mathematical Modeling of Biological Systems, Heinrich-Heine University of Düsseldorf, Germany

## Abstract

The ability to design RNA molecules with specific structures and functions could facilitate research and developments in biotechnology, biology and pharmacy. Here we present a flexible RNA design framework based on deep learning that locally optimizes sequences by gradient-guided search methods. We demonstrate its effectiveness by designing bi-stable RNA molecules by superimposing conformer target structures.

## 1 Introduction

There is a need for algorithms that can reliably generate RNA sequences with desired structures and functions. Ranging from biosensors that report metabolite concentrations in vivo, ^1^ to translationally optimized transgenes^2^ or highly stable messenger RNA molecules relevant for RNA therapeutics, ^3^ designing RNA molecules with specific functions remains a challenge in biotechnology, biological and pharmaceutical research.

Recent advances in deep learning that have enabled accurate protein design^4^ suggest that similar algorithms could also enable RNA design. However, training of the involved deep neural networks requires vast amounts of structural data, of which there exists far less for RNA molecules than for proteins. Instead of designing RNA molecules with specific 3D structures, RNA design has so far been limited to the design of RNA molecules with specific secondary structures.^5,6^ RNA secondary structure describes all base pairing interactions within one or between several RNA molecules and they can be readily represented by adjancency matrices called contact maps.^7,8^ This has been shown to be sufficient for the design of certain functional RNA molecules including riboswitches.^9–11^ These functional RNA molecules have in common that they are multi-stable: unlike proteins, which typically fold into one structure, RNA molecules can fold into several stable conformations with frequencies given by the Boltzmann-distribution (conformers with higher melting temperatures occur more frequently). Numerous algorithms have been developed that can accurately and rapidly predict secondary structures, a requirement for RNA design (see Supplementary Information). A special secondary structure pattern called pseudoknots, where residues of a loop interact with residues outside the loop, can only be predicted by some algorithms as their prediction drastically increases the complexity.^7,12^

*’* A major challenge arises when attempting to compare RNA design methods. Although X-ray crystallography and NMR have become indispensable methods in structural biology, the involved work-flows can be labour-intensive and expensive. Moreover, while chemical probing methods such as SHAPE^13^ or PARS^14^ can give clues about which residues are likely engaging in base pairing interactions, these methods cannot directly reveal the secondary structure. Nondenaturing polyacrylamide gel electrophoresis (PAGE), on the other hand, is a straight-forward method that can give insight into certain global structural properties of nucleic acid molecules.^15^ For example, nucleic acids with slim structures migrate faster through gels than bulky ones, because the latter get entangled between the gel polymers more easily.^16–18^ This effect has been used to separate RNA conformations from conformational mixtures such as *in vitro* transcribed RNA.^19^

Here we investigated the question of whether multi-stable RNA molecules could be designed with deep learning. We hypothesized that a deep neural network that was trained to predict secondary structures from sequence information could be used to optimize RNA sequences so that the predicted secondary structures matched the structures of the desired conformers. We tested the framework’s functionality by designing bi-stable RNA molecules with a slim and a bulky conformation and confirmed the design’s success with non-denaturing PAGE.

## 2 MATERIALS AND METHODS

### 2.1 Algorithm

A detailed description of the algorithm is provided in the Supplementary Information. In short, we first created a secondary structure model that could predict contact maps for input sequences. The model and trained on a synthetic data set comprised of 1 million random sequences and corresponding contact maps that were predicted by RNAfold.^20^ Then, the secondary structure model was run ”in reverse”: a target contact map describing the desired secondary structure was provided, and a sequence variable was iteratively adjusted so that its predicted contact map became as close to the target contact map as possible, which was done by gradient descent or discrete optimization. To design bi-stable RNA molecules, the superposition (average) of the individual conformer contact maps was used as target.

### 2.2 Reagents

#### 2.2.1 Constructs

Template DNA constructs for *in vitro* transcription consisted of the *T*7 promoter, the target gene and the HDV ribozyme to promote sharp bands ^21,22^ (see Supplementary Information for detailed sequence information). Three additional guanines were inserted between the T7 promoter and the target gene to enhance transcription.^23^ All constructs were obtained as vectors (MiniGene 25 – 500 bp in pIDTSMART-AMP from Integrated DNA Technologies GmbH, Germany) and were verified by Sanger sequencing.

### 2.3 RNA preparation

Template DNA was amplified by PCR (see Supplementary Information for primers) and purified using gel excision (NucleoSpin Gel and PCR Clean-up kit, MACHEREY-NAGEL GmbH & Co. KG, Germany). Amplicons were transcribed in vitro for 24 hours at 37°C (HiScribe T7 Quick High Yield RNA Synthesis Kit, New England Biolabs GmbH, Germany). RNA was purified by phenol-chloroform extraction (miRNeasy Mini kit, QIAGEN GmbH, Germany) with 700 μl QIAzol lysis reagent per sample. To lift potential kinetic traps, RNA was melted at 70°C for 10 minutes in 10 mM MgCl_2_, followed by a cool-down to 4°C at 0.1°C × s^-1^.

### 2.4 Non-denaturing polyacrylamide gel electrophoresis (PAGE)

0.5× TBE Polyacrylamide gels (12% polyacrylamide) were prepared with 37.5 : 1 polyacrylamide / bisacrylamide solution. Of each RNA sample, 1 μl (1 μg × μl^-1^ RNA) was loaded and gels were run at 16 V × cm^-1^ for 50 minutes (Mini-PROTEAN system, Bio-Rad Laboratories GmbH, Germany). Gels were stained with SYBR Green II for 15 minutes (Thermo Fisher Scientific GmbH, Germany).

### 2.5 Secondary structure prediction with third-party programs

To generate contact maps representing structural ensembles with RNAsubopt, ^20^ RNAsubopt was run at 37°C with the ”stochBT” option to randomly draw 10, 000 structures according to their ensemble probabilities. The output dot-bracket representations were individually converted to contact maps and averaged. Pseudoknots were predicted using the IPknot Webserver (URL: http://rtips.dna.bio.keio.ac.jp/ipknot/) with the default settings.^24^

### 2.6 Statistical Analyses

The similarity between two sequences *x* and *y* was statistically tested with a binomial distribution 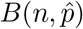 with *n* = 100 corresponding to the sequence length and matching probability estimated by 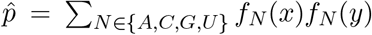, where *f_N_* denotes the frequency of nucleotide *N*.

### 2.7 Metrics

AUROC values relating predicted and target contact maps were calculated with thresholds restricted to values larger than 0.01 to avoid taking very small contact probabilities into account. Predictions were carried out for sequences with leading triple-guanine (GGG) to match the actual transcript sequences.

## 3 RESULTS

### 3.1 Design of bi-stable RNA-molecules

To test our RNA design algorithm, we chose to design bi-stable RNA molecules with two highly different conformations: a slim and a bulky structure, which we expected to migrate faster and slower through polyacrylamide gels, respectively (Figure 1A). The target secondary structures were designed by hand with an equal number of base pairs to avoid biased folding energies. We decided to design sufficiently long RNA molecules (100 nucleotides) to make rapid conformational switching negligible at 37°C so that sharp bands could be obtained with non-denaturing PAGE. We then carried out an *in silico* sequence search, which resulted in a significantly shifted loss distribution between initial and locally optimized sequences (Figure 1B). We selected the top-3 locally optimized bi-stable candidate sequences (Supplementary Information). Candidates 1 and 2 exhibited significantly greater sequence similarity (37, *p* = 0.02) than expected by chance.

**Figure 1:**
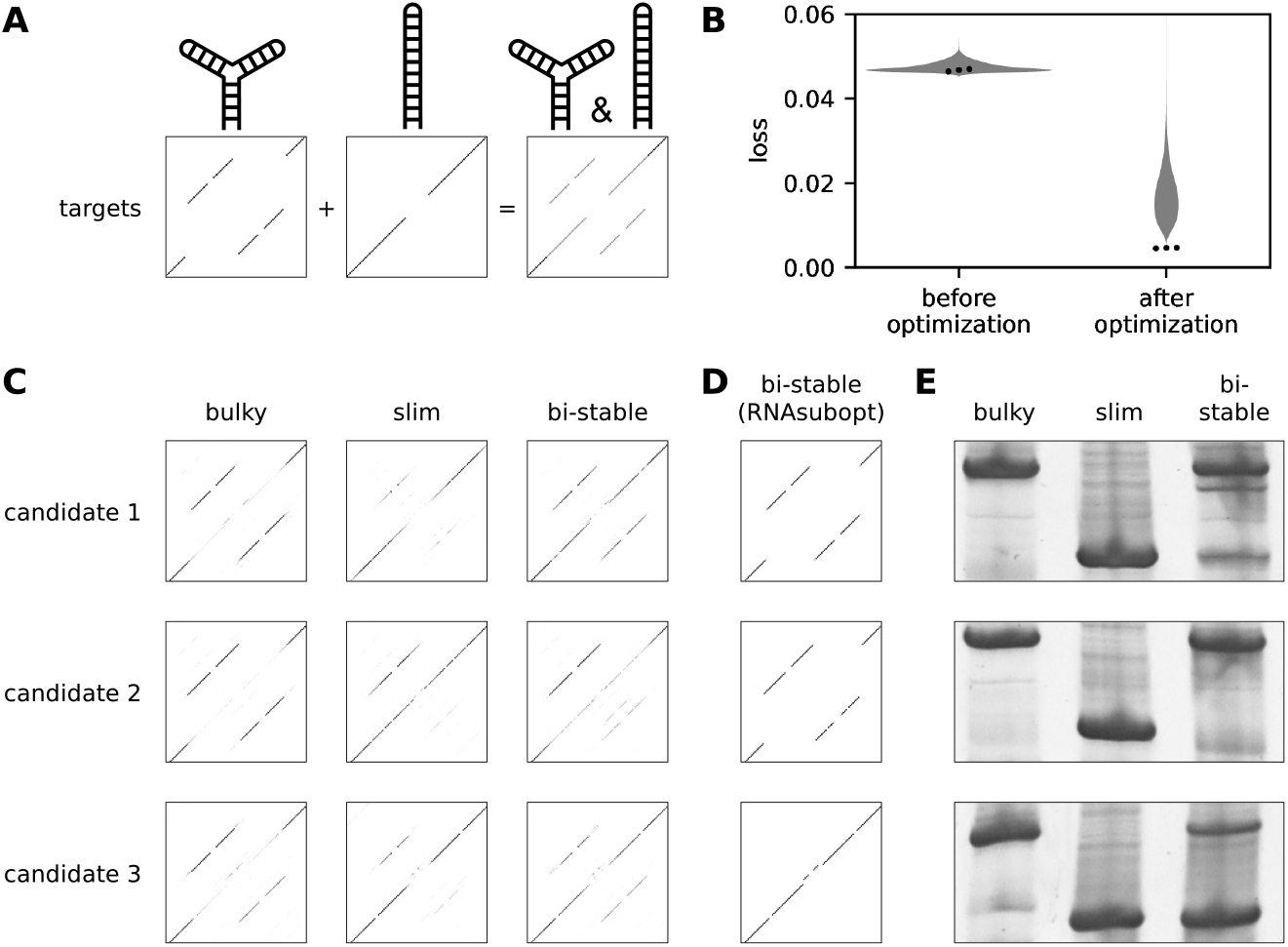
Design of bi-stable RNA molecules. **(A)** Target contact maps for the bulky, slim and bi-stable structures. The superposition of the contact maps for the bulky and slim structures (left and center, respectively) results in the contact map for the bi-stable structure (right). **(B)** Violin plot of the loss values between predicted and target structures before and after gradientdescent-based sequence improvement during random sequence search. The loss values of the top-3 candidates are indicated by the three dots (left to right). **(C)** Predicted contact maps of the bi-stable candidates (right) and their bulky and slim mutants (left and center, respectively); contact maps for candidates 1 – 3 are at the top, middle and bottom, respectively. **(D)** Contact maps of the top-3 candidates as predicted by RNAsubopt. **(E)** Non-denaturing PAGE of RNA molecules for the top-3 bi-stable candidates and their respective slim and bulky mutants.

Before analysing candidate RNA molecules with non-denaturing PAGE, we sought for a way to identify which bands belonged to which conformer. We reasoned that mutating the bi-stable candidates at the right sites could cause the resulting RNA molecules to fold into only either the slim or bulky conformation, and that those molecules would migrate with approximately the same speeds as the conformers of the respective bi-stable candidates. For each bi-stable candidate, we created two mutants by discrete sequence optimization that were predicted to fold into only the slim or bulky conformation. Moreover, the mutants differed from the respective bi-stable candidates by two point mutations. The predicted contact maps of the bi-stable and mutant candidates were highly similar to their targets (Figure 1C, Table 1), and bi-stable candidates also displayed substantial similarity to both slim and bulky targets. In contrast, RNAsubopt did not predict any candidate to be bi-stable, with predicted contact maps having lower similarity to the bi-stable target but high similarity to either the bulky or slim targets (Figure 1D, Table 1). Next, we prepared RNA for all bi-stable and mutant candidates and analysed RNA structures by non-denaturing PAGE. The first and third bi-stable candidates exhibited two dominant bands indicating the presence of two separable conformations (Figure 1E). The second bi-stable candidate exhibited one dominant and one faint band whose presence was confirmed by 2 replicate experiments (Supplementary Figure S3). The first bi-stable candidate also exhibited a weak satellite band that migrated slightly faster than the slowly migrating band. This indicated the presence of another, rarely assumed conformation, which was not predicted by either the secondary structure model or RNAsubopt. All slim and bulky mutant candidates exhibited one dominant band. In addition to the dominant band, the bulky mutant of the third bi-stable candidate also displayed a faint, faster migrating band, indicating an incomplete equilibrium shift. The dominant bands of all slim mutants migrated faster than their bulky counterparts. Moreover, the dominant bands of the bi-stable candidates migrated comparably fast as their respective slim and bulky mutants. Taken together, these observations indicate that the second and third bi-stable candidates were indeed bi-stable and that the conformers corresponded to the slim and bulky conformations. While the first bi-stable candidate appeared to be tri-stable instead of bi-stable, two bands migrated with similar speeds as the respective slim and bulky mutants, indicating that also in this case, two conformers of the bi-stable candidate corresponded to the slim and bulky conformations.

**Table 1.** AUROC values between target and predicted contact maps. All values were calculated for sequences with leading triple-guanine (GGG).

| prediction method | candidate | variant | Target |  |  |
| --- | --- | --- | --- | --- | --- |
|  |  |  | bulky | slim | bi-stable |
| Secondary structure model | 1 | bi-stable | 1.00 | 1.00 | 1.00 |
|  |  | slim | 0.88 | 1.00 | 0.93 |
|  |  | bulky | 1.00 | 0.92 | 0.96 |
|  | 2 | bi-stable | 0.99 | 0.96 | 0.97 |
|  |  | slim | 0.91 | 0.97 | 0.94 |
|  |  | bulky | 1.00 | 0.80 | 0.88 |
|  | 3 | bi-stable | 0.98 | 0.98 | 0.98 |
|  |  | slim | 0.84 | 0.98 | 0.89 |
|  |  | bulky | 1.00 | 0.96 | 0.98 |
| RNAsubopt | 1 | bi-stable | 1.00 | 0.69 | 0.81 |
|  |  | slim | 0.66 | 1.00 | 0.80 |
|  |  | bulky | 1.00 | 0.69 | 0.81 |
|  | 2 | bi-stable | 1.00 | 0.69 | 0.81 |
|  |  | slim | 0.66 | 1.00 | 0.80 |
|  |  | bulky | 1.00 | 0.69 | 0.81 |
|  | 3 | bi-stable | 0.66 | 0.98 | 0.79 |
|  |  | slim | 0.66 | 0.98 | 0.79 |
|  |  | bulky | 1.00 | 0.69 | 0.81 |

### 3.2 Design of pseudoknots

We sought to examine if the framework could also be used to design pseudoknots, even though the secondary structure model had not been trained on synthetic data containing pseudoknots. We argued that this should be possible nonetheless because pseudoknot contact maps can be decomposed into multiple, non-pseudoknot contact maps (Figure 2A). Because we could not design an experiment that could have revealed the presence of pseudoknots with non-denaturing PAGE and because the use of X-ray crystallography was beyond our resources, we resorted to analysing candidates *in silico* with IPknot, an independent program capable of predicting pseudoknots. ^24^ We designed a target pseudoknot secondary structure by hand (Figure 2A, Supplementary Information), carried out sequence search and selected the top-3 candidates (see Supplementary Information for candidate sequences). The second candidate was predicted to be a pseudoknot by both the model and IPknot (Figures 2B and 2C, respectively).

**Figure 2:**
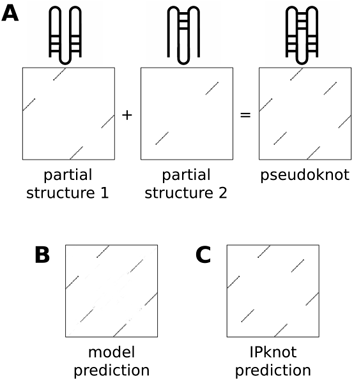
Design of pseudoknot RNA molecules. **(A)** The superposition of two partial contact maps results in a pseudoknot contact map. **(B)** Model contact map predction of the second pseudoknot candidate. **(C)** IPknot contact map predction of the second pseudoknot candidate.

## 4 DISCUSSION

### 4.1 Designing bi-stable structures with specific conformer frequencies is hard

While all candidates had at least two conformations, we observed that the frequencies of the slim and bulky conformations did not match the desired 50 : 50 ratio. There are two explanations for this observation: either the candidate sequences were not sufficiently optimized, or the secondary structure model was inaccurate. Increasing the number of initial random sequences during sequence search could result in candidates that match the ratio more closely. If this does not improve the ratio, there are several possible reasons why the secondary structure model might have been inaccurate. Precisely matching a 50 : 50 conformation ratio is difficult because the conformation ratio is highly sensitive to small folding energy deviations in this regime. In fact, changing a conformation ratio from 50 : 50 to 99 : 1 requires an energy difference of 11.8 kJ × mol^-1^ (2.8 kcal × mol^-1^) at 37°C, which is on the order of the enthalpy of a weak hydrogen bond. ^25,26^ Moreover, entropic effects can have can have significant impact when two conformations have almost identical folding enthalpies. Because the energy model underlying the Vienna RNA package has only limited ability to model entropic effects, it is reasonable to assume that the same is true for the secondary structure model since it was trained on synthetic data generated by RNAfold. However, discerning between inaccurate enthalpy calculations or entropic effects is not straight-forward. Inaccurate secondary structure predictions could also be the result of insufficient model capacity to learn the complex sequence-structure relationship from data. This could be tested by investigating alternative neural network architectures or self-supervised pre-training techniques. Moreover, training the model on synthetic data corresponding structural ensembles rather than the most stable structures could be a way to improving secondary structure predictions.

### 4.2 Bias towards bi-stability

While the secondary structure model correctly predicted all candidates to be bi-stable, RNA-subopt did not. This may seem surprising given the fact that the secondary structure model was trained on data generated by RNAfold, which uses the same energy function as RNAsubopt. Moreover, the experiments showed that all but one mutant were not bi-stable, but their contact map predictions still showed substantial similarity to the respective other target. We argue that both observations are the result of the secondary structure model being biased towards predicting bi-stable structures. The most likely reason for the bias is the underlying neural network’s limited ability to model mutually exclusive patterns: in situations where multiple secondary structures match a sequence well, deciding which structure to favor can be difficult. This results in a tendency to output linear interpolations of partial contact maps, which can manifest as a bias towards predicting bi-stable structures. This may also be the reason why, although the secondary structure model was trained on data that did not contain pseudoknots, the framework seems to be capable of designing pseudoknots, that is, examples that lie outside the training distribution. A thorough investigation of different neural network architectures may reveal the extent of the bias and whether its reduction comes with a reduced ability to design pseudoknots.

### 4.3 Framework improvements

The described framework was investigated in a basic form with the aim of establishing a proof of principle and setting a baseline. This is why there may be room for algorithmic improvements concerning (i) the speed of convergence during sequence search, (ii) the quality of the predictions, (iii) the types of design constraints and (iv) the types of structures that can be designed. First, sequence search convergence could be accelerated by including prior knowledge about the underlying problem, including drawing initial sequences from a distribution that is already informed by the gradient and making use of genetic algorithms that exploit the fact that similar sequences have similar structures. Sequence initialization and genetic algorithms have already been used by several RNA design algorithms. ^9,27^ Second, secondary structure predictions could be improved by utilising neural network architectures that optimally combine local and global sequence features for contact map prediction. In this work, a residual network architecture with self-attention was used, but architectures such as U-nets^28^ that spatially compress contact maps into high-dimensional representations might enable more efficient processing of global sequence information or vision transformers^29^ that purely rely on the attention mechanism. Third, there may be situations in which additional design constraints need to be set. For example, it can happen that only a partial target structure can clearly be specified or that some structural features are irrelevant. This could be accounted for by excluding the respective target structure regions from the loss. Another, more involved scenario is riboswitch design.^30^ Given an aptamer domain with known energy release upon ligand binding, riboswitch design could be realized by specifying relative conformer abundances at different energy levels, that is, at the bound and unbound states. This could be achieved by (i) enhancing the framework with a model that can predict both secondary structures and corresponding folding energies, (ii) using weighted target contact maps and (iii) keeping the parts of the sequence variable corresponding to the aptamer domain fixed. Finally, the framework could be extended to enable 3-D RNA design, which could be accomplished by exchanging the secondary structure model with one that predicts 3-D structures. However, specifying 3-D target structures by hand is challenging and may require the development of an interactive user interface that gives immediate feedback about which structures are possible. We have implemented the framework in a modular way in PyTorch^31^ to facilitate the implementation of improvements and new features.

### 4.4 RNA design benchmark

Although RNA design algorithms should ideally be tested experimentally if the designed sequence has the desired structure, this is rarely done in practice. The reason is that experimental structure elucidation methods are expensive and labor-intensive. X-ray crystallography often requires the identification of adequate crystallization conditions and sometimes even sequence alterations to enable crystallization. Liquid NMR on the other hand does not require crystallization, but it can only be used to reconstruct 3-D structures of short RNA molecules. As a consequence, RNA design algorithms are commonly compared *in silico*. This is problematic: because different RNA molecules can have the same secondary structure, it is generally impossible to define a ground truth sequence for a given secondary structure without defining additional constraints such as a particular folding energy. Consequently, studies that have evaluated RNA design methods *in silico* by trying to reconstruct sequences for given known RNA structures have reported moderate success rates.^9,27,32,33^ Because non-denaturing PAGE is a cheap and straight-forward experimental method that allows the clear separation of structually distinct conformers and even quantitative analyses of ensemble frequencies, we advocate for using this simple experimental design as a benchmark for the comparison of RNA design methods.

## 5 ACKNOWLEDGEMENTS

We thank Graeme L. Conn and Gerhard Steger for helpful discussions and Sarah Rose Richards for language editing. Computational services were provided by the High-Performance Computing Platform of the Heinrich-Heine University in Düsseldorf, Germany.

## 5.0.1 Funding

This work was supported by the European Regional Development Fund (ERDF) [grant number 0400380].

## Supplementary Information

### 1 Introduction

#### 1.1 RNA design

RNA design is an inverse problem as the goal is to infer a cause (the RNA sequence) from an effect or observable property (position of atoms in space), hence RNA design is also referred to as inverse folding. There are two major challenges in RNA secondary structure design. First, some desired secondary structures are physically impossible to realize by any sequence under reasonable conditions. In such a case, it is desirable to obtain a sequence with a similar but possible secondary structure. Second, the same secondary structure may be assumed by several sequences. This means that, in general, no direct map from the secondary structure space to the sequence space can be learned. In order to output a single solution, an algorithm must make at least one “decision” about which of the possible sequences to produce. These decisions can either be stochastic or deterministic. Most algorithms are stochastic and employ local or global search methods to identify sequences with a desired secondary structure. ^1^ The sequence space search is carried out by randomly sampling from a sequence distribution, and the search space can be restricted by heuristics including deep learning models. Typically, many sequences must be tested for sufficient structural similarity to the desired target, so this approach relies on the ability to efficiently compute secondary structures from sequence information. The advantage of stochastic search is that multiple sequences can be examined and the best fitting one can be chosen, but the sequence search may be computationally intensive. There are several stochastic methods that can design bi-stable RNA molecules by identifying sequences that can fold into two specified conformations, however these approaches rely on energy functions with empirically determined parameters that limit their flexibility. Deterministic sequence design, on the other hand, is realized by following a complex strategy to choose the right nucleotides in a given context. Deep neural networks have excelled at modelling complex relationships such as protein folding.^2^ Consequently, a recent deterministic algorithm is based on deep reinforcement learning that deterministically builds up sequences nucleotide by nucleotide after being trained on a large set of synthetic secondary structure and sequence pairs.^3^ While deterministic algorithms can be orders of magnitude faster than stochastic search methods, they only output one sequence and generally do not give information about how well a particular sequence fits to a secondary structure relative to other sequences.

#### 1.2 Secondary structure representations

RNA secondary structures describe all base pairing interactions within one or between several RNA molecules. Secondary structure representations of RNAs are sufficiently informative to enable modelling of multi-stable RNA structures, are straight-forward to interpret and have consequently been used in numerous RNA design approaches.^4^ Although numerous types of base pairing interactions have been classified, ^5^ Watson-Crick base pairs (A·G, C·U) and Wobble base pairs (G·U) are the most common types. This is why most secondary structure representations only represent those two interaction types. There are two common secondary structure representations. Contact maps are adjacency matrices that indicate which residues form base pairs with one another, and dot-bracket notation uses strings of balanced opening and closing brackets to indicate base pairs along sequences.^6^

### 2 Algorithm

#### 2.1 Framework components

The RNA design framework consisted of five main components: an RNA sequence variable *x*, a target secondary structure *y*, a differentiable structure prediction model *M*, a loss function *F* and an optimization routine (Figure S1).

The sequence variable *x* was either continuous or discrete, depending on the optimization routine (described below). The main purpose of the continuous representation was that it was required for sequence optimization with backpropagation. In both continuous and discrete cases, the sequence variable was an (*L* × 4) matrix, where *L* denotes the sequence length and 4 corresponds to the four possible nucleotide states. In the discrete case, x was one-hot encoded so that *x_i,k_* = 1 and *x_i,l≠k_* = 0 indicated that the i-th nucleotide of the sequence was the k-th nucleotide in the ordered set {*A*, *C*, *G*, *U*}. In the continuous case, *x_i,k_* ∈ [0,1], which was ensured by the normalization ∑*_k_ e^x_i,k_^* = 1.

The secondary structure model *M* took an input sequence *x* and produced an (*L* × *L*) contact map prediction *ŷ*, where *ŷ_i,j_* ∈ [0,1] denotes the predicted contact probability between residues *i* and *j*. The model was implemented as a deep residual network.^7^ A hard-coded transformation was used to convert input sequences into image-like (*L* × *L* × *D*) tensors that could be used by the network, with *D* = 16 corresponding to the number of possible nucleotide pairing combinations. The network consisted of 8 residual and 8 patch-wise self-attention layers that were used alternatingly (self-attention is a special mechanism that is at the core of recent large language models and AlphaFold^2^). The aim of patch-wise self-attention was to enable information exchange between non-local features at each layer.^8,9^ It was realized by dividing each layer into 8 equally-sized patches, after which self-attention was used to weight the patches (see section 2.2 for details). All residual and patch-wise self-attention layers had 8 channels. The secondary structure model was trained on 1 million one-hot encoded random sequences and corresponding contact maps that were predicted by RNAfold.^10^

Target secondary structures were encoded as (*L* × *L*) contact maps. To specify targets for bi-stable RNA molecules with individual conformer contact maps *y*_1_ and *y*_2_, the average *y* = 0.5 × (*y*_1_ + *y*_2_) was used.

The binary cross-entropy (log-likelihood of the Bernoulli distribution) over all predicted base pairs served as the loss function between target and predicted contact maps:

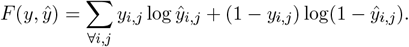

Sections 2.3 and 2.4 describe the details of the search routine, which was either based on gradient descent that could change multiple nucleotides per optimization step, or by discrete optimization that changed a specified number of nucleotides per step.

**Figure S1:**
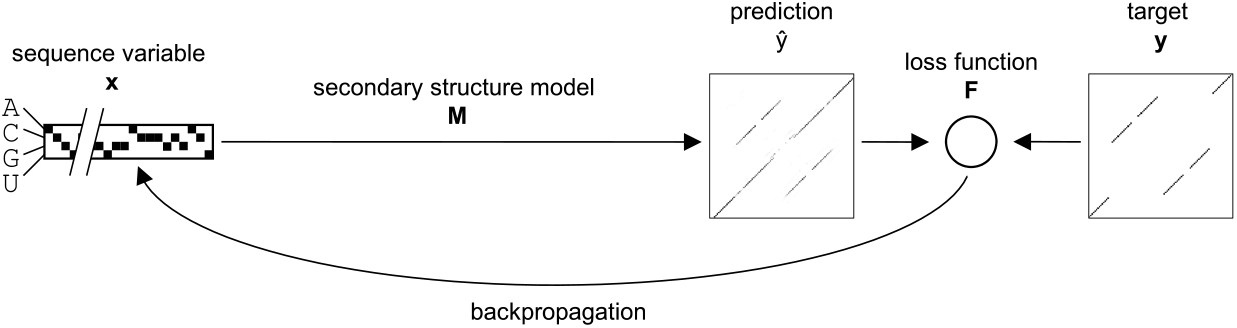
Framework components (except search-routine).

#### 2.2 Patch-wise self-attention

Input to each patch-attention layer was an (*L* × *L* × *D*) tensor *z^in^*, with *L* denoting the width and height, and *D* the number of channels (Figure S2). We then applied convolution with kernel size *k* and stride *s* = *k* to obtain a spatially compressed (*K* × *K* × *D*) tensor, where *K* = *L*/*k*. Because the network’s output should eventually result in predictions for contact maps, which are symmetric, we extracted the upper triangular matrix without the diagonal to obtain matrix *x* with shape (*H* × *D*), where *H* = *K* (*K* – 1)/2. We then applied self-attention. In simple terms, we calculated attention scores based on learnable spatial contact correlations. Following a commonly used notation on non-local neural networks, ^9^ the layer output is given by:

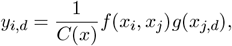

where 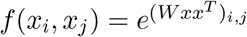, *C*(*x*) = ∑_*j*_*f*(*x_i_*,*x_j_*), *g*(*x_i,d_* = *x_i,d_*, and *W* is an (*H* × *H*) matrix. We then used all *y_i,d_* to fill the upper and lower triangular part of a (*K* × *K* × *D*) matrix *r*, which was blown up in size to (*L* × *L* × *D*) and added to the original input tensor *z* to yield output tensor *z^out^*, according to: 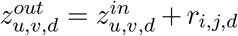, where *i* = ⌈*u*/*k*⌉ and *j* =⌈*v*/*k*⌉ *k* is the kernel size).

**Figure S2:**
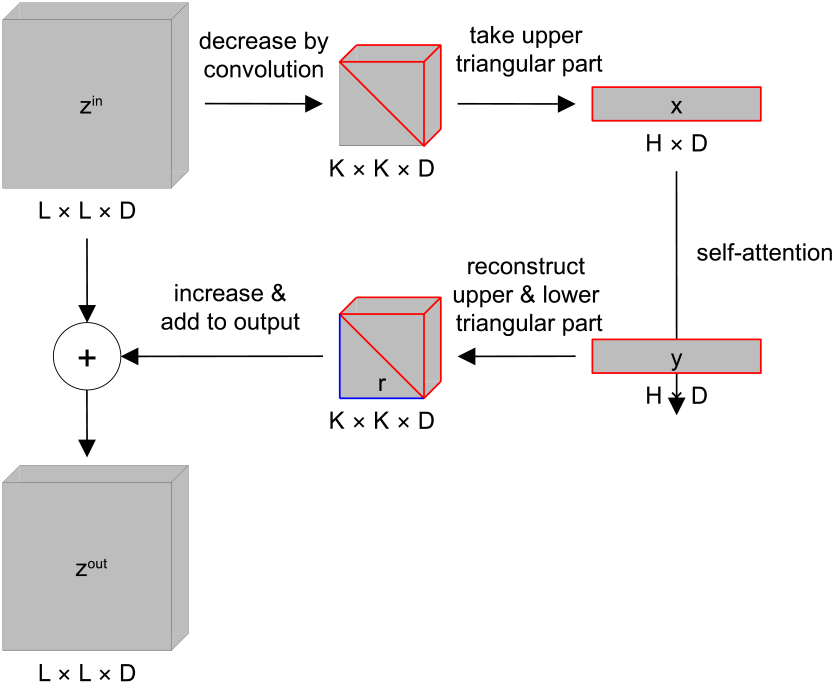
Patch-wise self-attention.

#### 2.3 Candidate sequence search

The candidate sequence search routine was a random search combined with gradient descent. First, a large number of random, unbiased sequences were drawn that served as independent initializations of the continuous sequence variable *x.* Then, for each initialization, the sequence variable was optimized by gradient descent so that the loss between the predicted and target secondary structures was minimized to a local optimum. At each gradient descent step *t*, the sequence variable was updated using the gradient of the loss between the target and predicted secondary structures according to: *x*_*t*+1_ ← *x_t_* + λ▽_*x*_*F*(*y*, *M*(*x_t_*)), followed by a normalization of the sequence variable so that ∑_*k*_*e^x_i,k_^* = 1. Upon convergence, the sequence variable was one-hot encoded. We used 10 million independent initializations and carried out 50 gradient descent steps per initialization.

#### 2.4 Mutant sequence search

In order to control the number of mutations at each optimization step, a discrete, one-hot encoded sequence variable *x* was used. The gradient of the loss between the target and predicted secondary structures, ▽*_x_F*(*y*,*M*(*x*)), was used to choose the mutations with the largest associated loss reduction.

### 3 Experimental details

#### 3.1 Primers

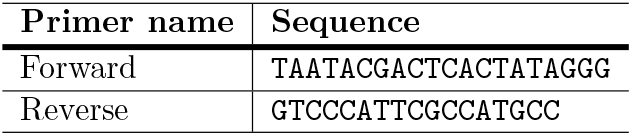

#### 3.2 General construct sequence

In the following example, the sequence of the first bi-stable candidate was used as target gene.

Color coding: T7 promoter / <u>extra guanines</u> / target gene / HDV ribozyme

~~~
TAATACGACTCACTATA<u>GGG</u>TCCGAATGCACCACTTTTAGGTCTCGTTTGTTTACGGGCGGGGCCTAACATTTCG
CTCTCTGCTCGTGGACGGGTGGGGGGTGGGGGTGGTGCGTTCGGAGCTAGCCATGGTCCCAGCCTCCTCGCTGGC
GGCTAGTGGGCAACATGCTTCGGCATGGCGAATGGGAC
~~~

### 4 RNA Design

#### 4.1 Bi-stable RNAs

##### 4.1.1 Target secondary structures

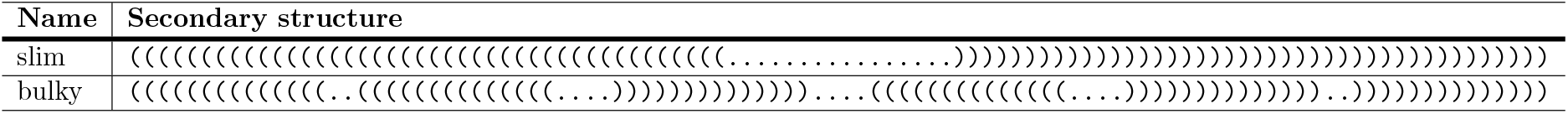

##### 4.1.2 Candidate sequences

Points mutations are <u>highlighted</u>.

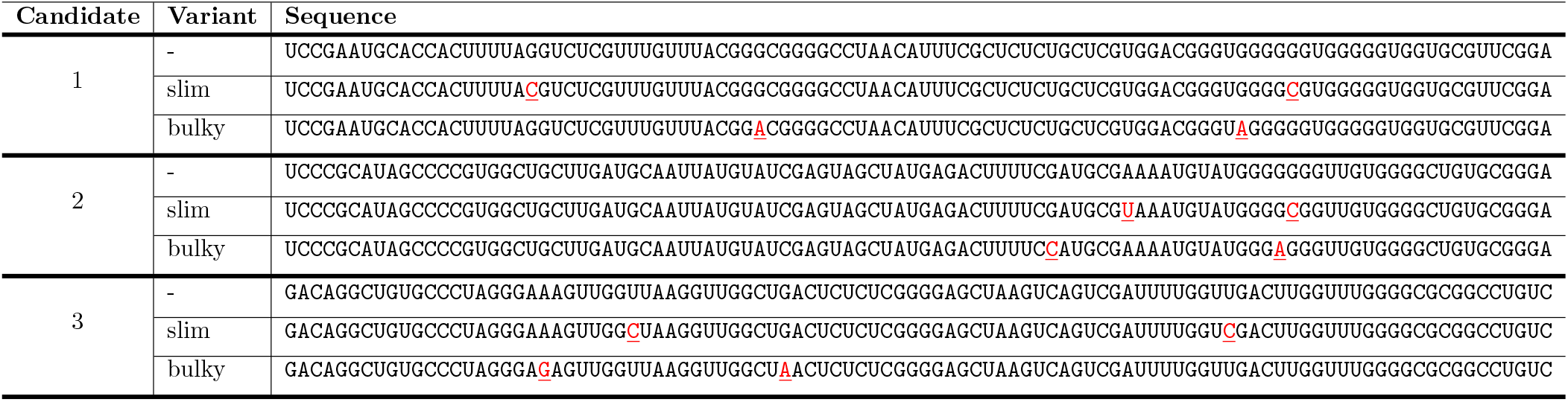

#### 4.2 Pseudoknot

##### 4.2.1 Target secondary structures

~~~
((((((((((((((..[[[[[[[[[[[[[[....))))))))))))))....((((((((((((((....]]]]]]]]]]]]]]..))))))))))))))
~~~

##### 4.2.2 Candidate sequences

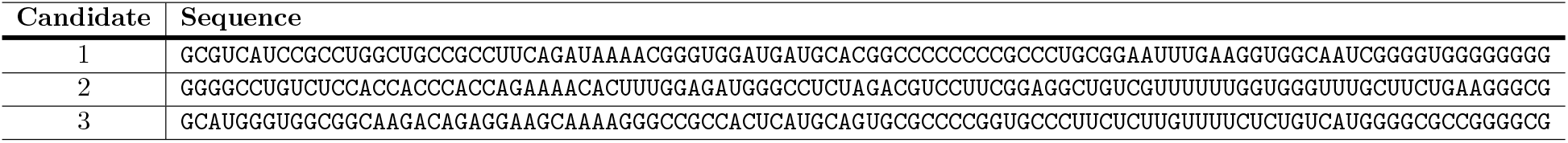

##### 4.2.3 Predicted secondary structures

Secondary structure predictions were carried out with IPknot using the default settings.

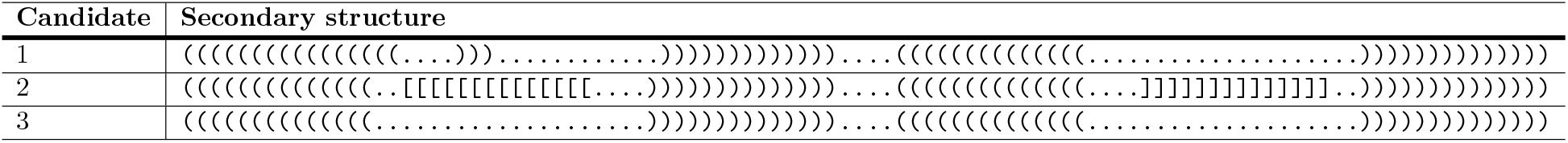

### 5 Replicate experiments

**Figure S3:**
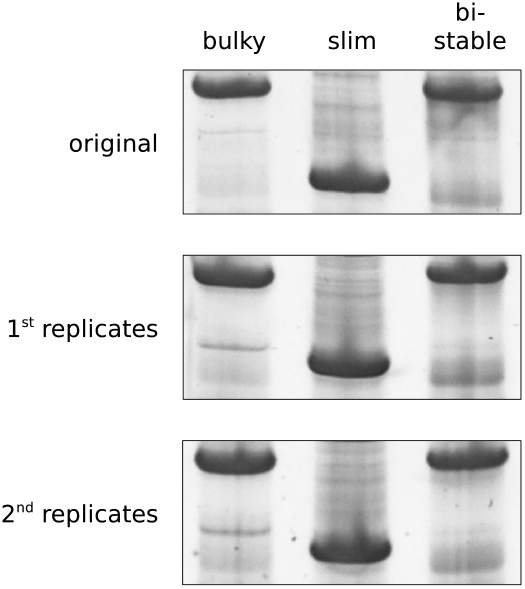
Replicate experiments: non-denaturing PAGE of RNA molecules for the second bi-stable candidate and the respective bulky and slim mutants. The original gel image is also shown in the main part of this publication.

## Notes

### Competing Interest Statement

The authors have declared no competing interest.

## References

[1] Hallberg, Z. F.; Su, Y.; Kitto, R. Z.; Hammond, M. C. https://doi.org/10.1146/annurev-biochem-060815-014628 2017, 86, 515–539.

[2] Verma, M.; Choi, J.; Cottrell, K. A.; Lavagnino, Z.; Thomas, E. N.; Pavlovic-Djuranovic, S.; Szczesny, P.; Piston, D. W.; Zaher, H. S.; Puglisi, J. D.; Djuranovic, S. Nature Communications 2019, 10.

[3] Wayment-Steele, H. K.; Kim, D. S.; Choe, C. A.; Nicol, J. J.; Wellington-Oguri, R.; Watkins, A. M.; Parra Sperberg, R. A.; Huang, P. S.; Participants, E.; Das, R. Nucleic Acids Research 2021, 49, 10604–10617.

[4] Jumper, J. et al. Nature 2021 596:7873 2021, 596, 583–589.

[5] Churkin, A.; Retwitzer, M. D.; Reinharz, V.; Ponty, Y.; Waldispühl, J.; Barash, D. Briefings in Bioinformatics 2018, 19, 350–358.

[6] Runge, F.; Stoll, D.; Falkner, S.; Hutter, F. 7th International Conference on Learning Representations, ICLR 2019 2019, 1–29.

[7] Hofacker, I. L.; Fontana, W.; Stadler, P. F.; Bonhoeffer, L. S.; Tacker, M.; Schuster, P. Monatshefte für Chemie / Chemical Monthly 1994 1 25:2 1994, 125, 167–188.

[8] Lu, X. J.; Bussemaker, H. J.; Olson, W. K. Nucleic Acids Research 2015, 43, e142–e142.

[9] Taneda, A. Frontiers in Genetics 2012, 3, 36.

[10] Findeiß, S.; Etzel, M.; Will, S.; Mörl, M.; Stadler, P. F. Sensors 2017, Vol. 17, Page 1990 2017, 17, 1990.

[11] Espah Borujeni, A.; Mishler, D. M.; Wang, J.; Huso, W.; Salis, H. M. Nucleic Acids Research 2016, 44, 1.

[12] Lyngsø, R. B.; Pedersen, C. N. S. Proceedings of the fourth annual international conference on Computational molecular biology - RECOMB ’00 2000,

[13] Cordero, P.; Lucks, J. B.; Das, R. Bioinformatics 2012, 28, 3006–3008.

[14] Kertesz, M.; Wan, Y.; Mazor, E.; Rinn, J. L.; Nutter, R. C.; Chang, H. Y.; Segal, E. Nature 2010, 467, 103–107.

[15] McPhie, P.; Hounsell, J.; Gratzer, W. B. Biochemistry 1966, 5, 988–993.

[16] Yuan, C.; Rhoades, E.; Heuer, D. M.; Saha, S.; Lou, X. W.; Archer, L. A. Analytical Chemistry 2006, 78, 6179–6185.

[17] Lilley, D. M.; Bhattacharyya, A.; McAteer, S. http://dx.doi.org/10.1080/02648725.1992.10647893 2013, 10, 379–401.

[18] Woodson, S. A.; Koculi, E. Methods in enzymology 2009, 469, 189.

[19] Calderon, B. M.; Conn, G. L. RNA 2017, 23, 557–566.

[20] Lorenz, R.; Bernhart, S. H.; Höner zu Siederdissen, C.; Tafer, H.; Flamm, C.; Stadler, P. F.; Hofacker, I. L. Algorithms for Molecular Biology : AMB 2011, 6, 26.

[21] Tanner, N. K.; Schaff, S.; Thill, G.; Petit-Koskas, E.; Crain-Denoyelle, A. M.; Westhof, E. Current Biology 1994, 4, 488–498.

[22] Schürer, H.; Lang, K.; Schuster, J.; Mörl, M. Nucleic acids research 2002, 30.

[23] Conrad, T.; Plumbom, I.; Alcobendas, M.; Vidal, R.; Sauer, S. Communications Biology 2020 3:1 2020, 3, 1–8.

[24] Sato, K.; Kato, Y.; Hamada, M.; Akutsu, T.; Asai, K. Bioinformatics 2011, 27, i85–i93.

[25] Serra, M. J.; Turner, D. H. Methods in Enzymology 1995, 259, 242–261.

[26] Halder, A.; Data, D.; Seelam, P. P.; Bhattacharyya, D.; Mitra, A. ACS Omega 2019, 4, 7354–7368.

[27] Lyngsø, R. B.; Anderson, J. W.; Sizikova, E.; Badugu, A.; Hyland, T.; Hein, J. BMC Bioinformatics 2012, 13, 1–12.

[28] Fu, L.; Cao, Y.; Wu, J.; Peng, Q.; Nie, Q.; Xie, X. Nucleic Acids Research 2022, 50, e14.

[29] Dosovitskiy, A.; Beyer, L.; Kolesnikov, A.; Weissenborn, D.; Zhai, X.; Unterthiner, T.; Dehghani, M.; Minderer, M.; Heigold, G.; Gelly, S.; Uszkoreit, J.; Houlsby, N. 2020,

[30] Domin, G.; Findeiß, S.; Wachsmuth, M.; Will, S.; Stadler, P. F.; Mörl, M. Nucleic Acids Research 2017, 45, 4108.

[31] Paszke, A. et al. PyTorch: An Imperative Style, High-Performance Deep Learning Library. Advances in Neural Information Processing Systems. 2019,.

[32] Andronescu, M.; Fejes, A. P.; Hutter, F.; Hoos, H. H.; Condon, A. Journal of Molecular Biology 2004, 336, 607–624.

[33] Busch, A.; Backofen, R. Bioinformatics 2006, 22, 1823–1831.

## References

[1] Churkin, A.; Retwitzer, M. D.; Reinharz, V.; Ponty, Y.; Waldispühl, J.; Barash, D. Briefings in Bioinformatics 2018, 19, 350–358.

[2] Jumper, J. et al. Nature 2021 596:7873 2021, 596, 583–589.

[3] Runge, F.; Stoll, D.; Falkner, S.; Hutter, F. 7th International Conference on Learning Representations, ICLR 2019 2019, 1–29.

[4] Churkin, A.; Retwitzer, M. D.; Reinharz, V.; Ponty, Y.; Waldispühl, J.; Barash, D. Briefings in Bioinformatics 2018, 19, 350-358.

[5] Lu, X. J.; Bussemaker, H. J.; Olson, W. K. Nucleic Acids Research 2015, 43, e142–e142.

[6] Hofacker, I. L.; Fontana, W.; Stadler, P. F.; Bonhoeffer, L. S.; Tacker, M.; Schuster, P. Monatshefte für Chemie / Chemical Monthly 1994 1 25:2 1994, 125, 167–188.

[7] He, K.; Zhang, X.; Ren, S.; Sun, J. Proceedings of the IEEE Computer Society Conference on Computer Vision and Pattern Recognition 2016, 2016-Decem, 770–778.

[8] Vaswani, A.; Shazeer, N.; Parmar, N.; Uszkoreit, J.; Jones, L.; Gomez, A. N.; Kaiser, L.; Polosukhin, I. Advances in Neural Information Processing Systems 2017, 2017-Decem, 5999–6009.

[9] Wang, X.; Girshick, R.; Gupta, A.; He, K. Proceedings of the IEEE Computer Society Conference on Computer Vision and Pattern Recognition 2018, 7794–7803.

[10] Lorenz, R.; Bernhart, S. H.; Höner zu Siederdissen, C.; Tafer, H.; Flamm, C.; Stadler, P. F.; Hofacker, I. L. Algorithms for Molecular Biology : AMB 2011, 6, 26.

